# APOBEC mutagenesis is low in most types of non-B DNA structures, unlike other types of cancer mutagenesis

**DOI:** 10.1101/2021.10.12.464135

**Authors:** Gennady V. Ponomarev, Bulat Fatykhov, Vladimir A. Nazarov, Ruslan Abasov, Evgeny Shvarov, Nina-Vicky Landik, Alexandra A. Denisova, Almira A. Chervova, Mikhail S. Gelfand, Marat D. Kazanov

**Affiliations:** Institute for Information Transmission Problems (the Kharkevich Institute, RAS), Moscow, Russia; Moscow Institute of Physics and Technology, Moscow, Russia; Dmitry Rogachev National Medical Research Center of Pediatric Hematology, Oncology and Immunology, Moscow, Russia; InterSystems Corporation, Cambridge MA, USA; Lomonosov Moscow State University, Moscow, Russia; Institut Pasteur, Paris, France; Skolkovo Institute of Science and Technology, Moscow, Russia

## Abstract

While somatic mutations are known to be enriched in genome regions with non-canonical DNA secondary structure, the impact of particular mutagens still needs to be elucidated. Here, we demonstrate that in human cancers, the APOBEC mutagenesis is not enriched in direct repeats, mirror repeats, short tandem repeats, and G-quadruplexes, and even decreased below its level in B-DNA for cancer samples with very high APOBEC activity. In contrast, we observe that the APOBEC-induced mutational density is positively associated with APOBEC activity in inverted repeats (cruciform structures), where the impact of cytosine at the 3’-end of the hairpin loop is substantial. Surprisingly, the APOBEC-signature mutation density per TC motif in the single-stranded DNA of a G-quadruplex (G4) is lower than in the four-stranded part of G4 and in B-DNA. The APOBEC mutagenesis, as well as the UV-mutagenesis in melanoma samples are absent in Z-DNA regions, due to depletion of their mutational signature motifs.

## Introduction

Recent studies of human cancer genomes revealed a significant role of the APOBEC (Apolipoprotein B mRNA Editing Catalytic polypeptide-like) family cytidine deaminases in cancer mutagenesis^1–3^. The APOBEC-family enzymes are a part of the human immune system acting against viruses and transposable elements^4^. APOBEC cytidine deaminases change the substrate DNA by binding to single-stranded DNA regions and deaminating cytosines in the TpC context, leading to C→T and C→G substitutions^5^. Positional clusters of somatic mutations with the APOBEC mutational signature were found in many types of cancers, in particular, breast, lung, bladder, head/neck, and cervical cancers^6–8^. It has been suggested that these mutation clusters arise when APOBEC binds to and slides along single-stranded DNA (ssDNA) accessible during replication, transcription, or double-strand breaks^9^. Recent studies support a link between these ssDNA-generated processes and APOBEC mutagenesis in the human cell^10^.

Several types of mutational heterogeneity along the genome, presumably associated with replication and transcription^11^, were identified for APOBEC mutagenesis. An increased density of APOBEC-induced mutations was found in early-replicating genome regions^12^, as opposed to other types of cancer mutagenesis, and in highly transcribed genes^13^. Higher rate of APOBEC-induced mutations was also observed on the lagging replicating DNA strand^14^ and the non-transcribed DNA strand^13^.

It is known that ssDNA, a preferred APOBEC substrate, can adopt diverse local conformations such as hairpins, loops, and pseudoknots^15^. A recent study has showed that cytosine at the 3’-end of a hairpin loop is a hotspot of APOBEC and can even be targeted by APOBEC while not being preceded by tymine^16^, essentially forming a second type of the APOBEC signature, dependent on the ssDNA secondary structure. Various forms of the ssDNA secondary structure have been found in the human genome regions with non-canonical DNA structure, that is, distinct from the right-handed DNA double-helix^17^.

To date, about ten types of non-B DNA structures are known, including hairpins/cruciform, triplexes (H-DNA), tetraplexes, slipped DNA, and Z-DNA. The cruciform structures are formed by inverted repeats^18,19^ that base-pair, forming an intrastrand hairpin stem and looping out the spacer between the repeat copies as ssDNA (Figure 1). Thus, the cruciform structure consists of two hairpin-loop arms and a four-way junction. The triplex DNA (H-DNA) structures can form at mirror repeats^20^, where ssDNA can bind in the major groove of the underlying DNA duplex forming a three-stranded helix (Figure 1). The four-stranded G-quadruplex structure is a co-planar array of four guanines formed by guanine-rich DNA stretches^21^ (Figure 1). The slipped strand DNA structures are formed when one strand of one copy of a direct repeat pairs with the complementary strand of another copy of a direct repeat^22^, yielding looped-out ssDNA (Figure 1). Sequences with an abundance of alternating purines and pyrimidines may form the double helix with a left-handed zigzag pattern called Z-DNA^23^ (Figure 1). As can be seen, most non-B DNA structures contain stretches of ssDNA, which might be expected to be an efficient substrate for the APOBEC enzymes.

**Figure 1.**
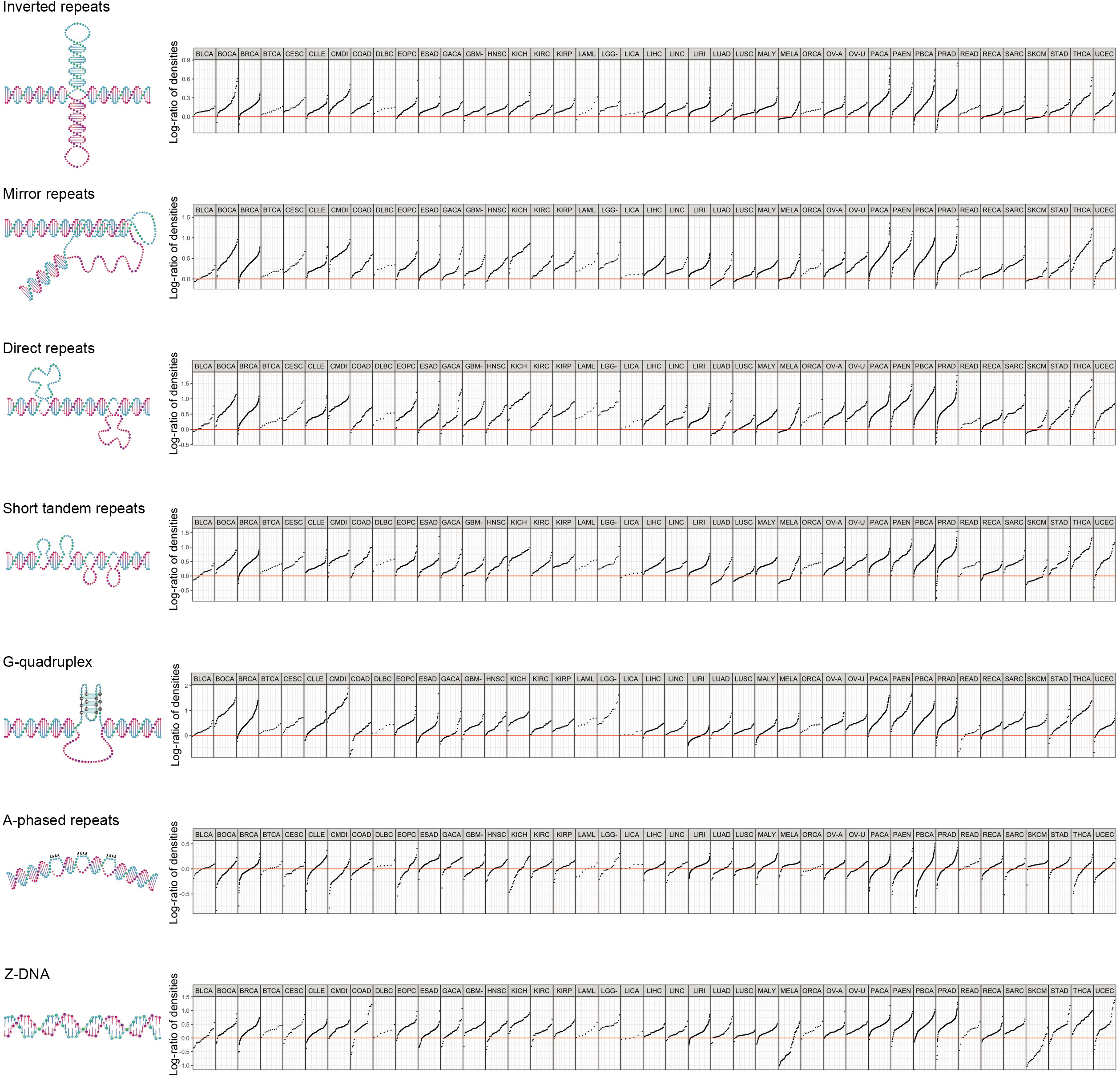
Enrichment of somatic single-base substitutions (SBS) observed in various types of non-canonical DNA structure genome regions and different cancers. Single dot corresponds to a particular cancer sample, with the vertical position indicating the log-ratio of the mutational density in non-B DNA structure genome regions to the mutational density in B-DNA genome regions.

Studies on cancer mutagenesis in non-B DNA genome regions showed an increased density of somatic mutations compared to the genome regions with conventual double-helix DNA structure^24^. Recent advances in the research on cancer mutagenesis have demonstrated that it is possible to assess the impact of particular mutagens using their mutational signatures. Here, we used the known mutational signature of APOBEC enzymes and analyzed the impact of the APOBEC-related mutagenesis in various types of non-B DNA genome structures, focusing on ssDNA regions. We found that, despite the presence of ssDNA in most non-B DNA structures, the APOBEC mutagenesis is not enriched in most non-B DNA structures, an exception being inverted repeats, and even is relatively lower in cancer samples with very high APOBEC activity.

## Results

### Single-base substitutions are generally enriched in non-B DNA structures in human cancers

To obtain a general view of somatic mutagenesis in non-B DNA structures in human cancer and use it further as a baseline for comparison with the APOBEC mutagenesis, we first calculated the densities of all somatic single-base substitutions (SBS) in non-B and B-DNA genome regions using mutational data from the Pan-Cancer Analysis of Whole Genomes (PCAWG) project^25^. Seven types of Non-B DNA structures were considered using genomic coordinates of non-B DNA motifs from the non-B DB^26^: direct repeats (DR), inverted repeats (IR), mirror repeats (MR), short tandem repeats (STR), G-quadruplexes (G4), A-phased repeats (APR), and Z-DNA. As was observed earlier for a smaller cancer dataset^24^, the density of somatic SBS is usually enriched in non-B DNA structures. Here, we used more than 1500 cancer samples from 41 cancer types and observed a similar enrichment with some exceptions (Figure 1). From the non-B DNA motif axis, the most prominent exception is that A-phased repeats are not associated with the enrichment of somatic SBS density: the mean fold enrichment among cancers was not statistically different from the zero enrichment (the Mann–Whitney–Wilcoxon test, *P*-value = 0.34). The same trend was observed earlier for germline mutations^27^. From the cancer axis, the most striking deviation was the decrease of mutational density in the Z-DNA motif for melanoma: by a factor of 2.16 (*P* = 9.1×10^−7^) for the MELA cancer type and by a factor of 3.06 (*P* = 4.8×10^−8^) for the SKCM cancer type. We have analyzed these deviations in more detail below (see the respective subsection). In all other cases, the mutational density was enriched in non-B DNA motifs: the mean of the mean fold enrichment over all cancer types was 3.0 for DR (*P* = 1.4×10^−14^), 2.7 for G4 (*P* = 7.1×10^−14^), 1.4 for IR (*P* = 1.4×10^−14^), 2.1 for MR (*P* = 1.4×10^−14^), 2.4 for STR (*P* = 1.4×10^−13^), and 1.9 for Z-DNA (*P* = 4.2×10^−9^). We also calculated enrichment in non-B DNA structures across all cancers separately for mutations attributed to the ABOPEC signature TpC motif (Supplemental Figure S1). The most noticeable difference compared to all mutations was the statistical insignificance of the enrichment (*P* = 0.06) in IR genome regions.

### APOBEC mutagenesis is not enriched in non-canonical DNA structures, unlike other types of mutagenesis, inverted repeats being an exception

To elucidate the characteristics of APOBEC mutagenesis in non-canonical DNA genome regions, we estimated the density of APOBEC-induced mutations for each type of non-canonical DNA structure in each cancer sample. We used the known APOBEC mutational signature TpC to extract mutations presumably associated with the APOBEC enzymes, excluding CpG-island regions, where APOBEC mutagenesis can overlap with hypermutation of unmethylated CpG sites. We first calculated the number of APOBEC-signature motifs, i.e., potential APOBEC targets, in each type of non-B DNA structure (Figure S2a). The fraction of TpC motifs for most types of non-B DNA structures was in the range of 0.07–0.12 except for Z-DNA, where this fraction was negligibly small, 0.008. Thus, we observed a depletion of the APOBEC-signature motif, TpC, in Z-DNA and have not further analyzed characteristics of the APOBEC mutagenesis in this type of non-B DNA structure.

For the remaining six types of non-canonical DNA structures, we calculated the density of APOBEC-induced mutations in cancer samples and compared it with the density of APOBEC-induced mutations in B-DNA genome regions in these samples. In four out of six non-B DNA motif types – DR, STR, MR, and G4 – we observed characteristics of APOBEC mutagenesis, which are different from the ones observed for other types of mutagenesis: the APOBEC-signature mutation density in non-canonical DNA genome regions was approximately equal to the density in B-DNA genome regions with the increase of APOBEC activity. Moreover, some cancer samples with a very high level of APOBEC mutagenesis demonstrated a lower density of APOBEC-signature mutations in non-canonical DNA genome regions compared to B-DNA genome regions.

More specifically, Figure 2a and Supplemental Figure S3 show how the log-ratio of the density of APOBEC-induced mutations within particular non-canonical DNA structure genome regions to the density of APOBEC-induced mutations in B-DNA genome regions depends on the activity of APOBEC mutagenesis in the samples. The activity of APOBEC mutagenesis for each cancer sample was estimated by the APOBEC enrichment as before^8^. It can be seen that samples with low APOBEC activity have an increased density of APOBEC-signature mutations in comparison with its density in B-DNA. This agrees with the fact that the density of somatic mutations in human cancer in non-canonical DNA motifs is generally higher that in canonical regions, as confirmed on PCAWG data in the previous section. It should be noted that most mutations with the APOBEC signature in these samples are likely associated with non-APOBEC mutagenesis, as the absence of APOBEC enrichment reflects a low level of APOBEC mutagenesis. Meanwhile, Figure 2a and Supplemental Figure S3 show that as the APOBEC activity increases, the APOBEC-signature mutation density in the four non-canonical DNA genome regions (DR, STR, MR, and G4) decreases to the level of the APOBEC-induced mutational density in B-DNA genome regions and to even lower level for cancer samples highly enriched in the APOBEC-signature mutations. The strongest effect was observed for the G4 structure.

**Figure 2.**
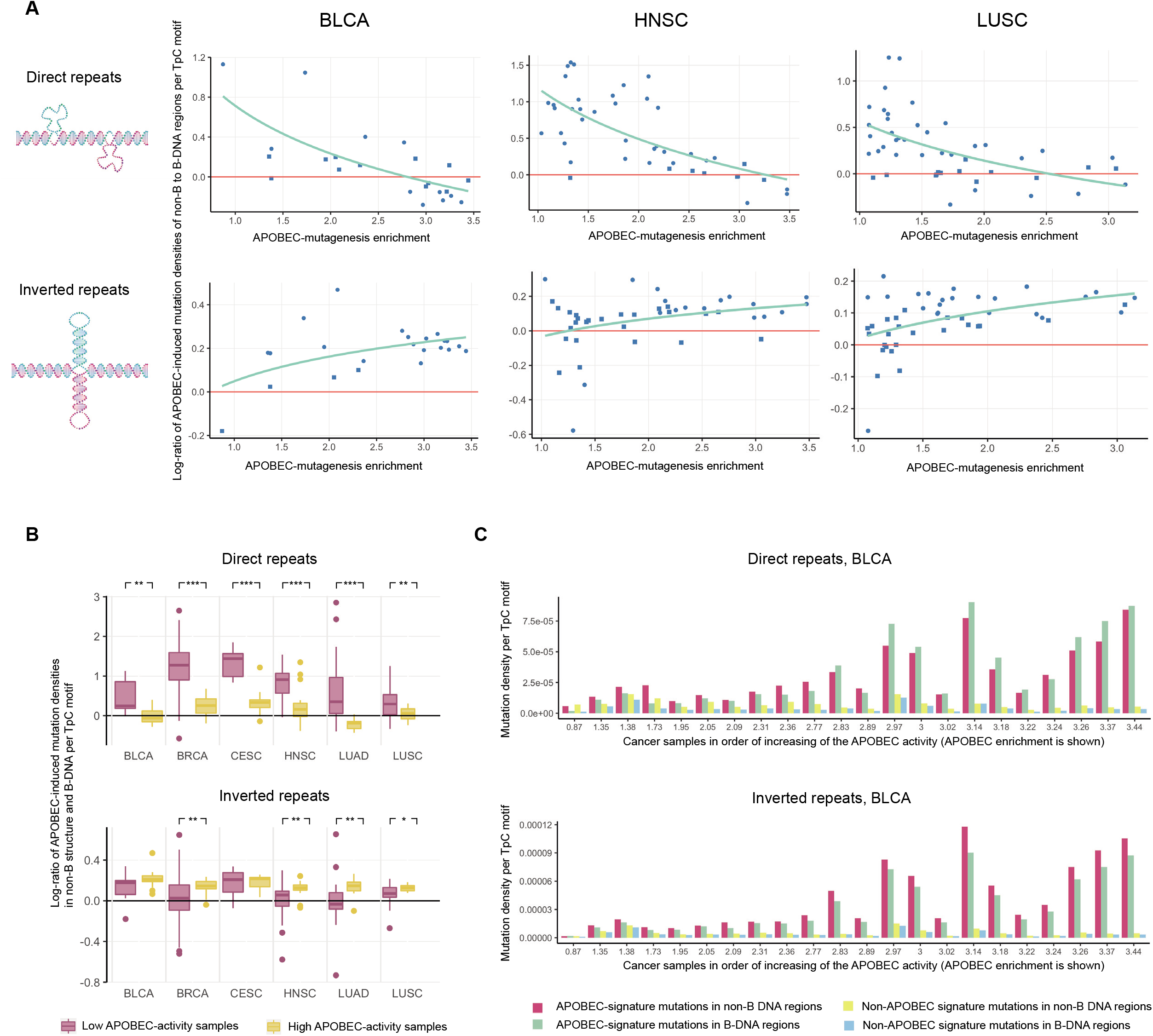
(a) Dependence of the log-ratio of APOBEC-induced mutation densities in non-B DNA to B-DNA genome regions on the activity of APOBEC mutagenesis in cancer samples. Point shape – round or square – corresponds to significant or insignificant statistical differences between two densities in a particular cancer sample, respectively. (b) Difference in distribution of the log-ratio of APOBEC-induced mutation densities in non-B DNA to B-DNA genome regions in cancer samples with low (APOBEC enrichment < 2.0) and high (APOBEC enrichment > 2.0) APOBEC activity. (3) APOBEC-induced and other mutation densities in non-B and B-DNA genome regions for bladder carcinoma (BLCA) samples. Notation: *** – *P*-value < 0.001, ** – *P*-value < 0.01, * – *P*-value < 0.05.

The behavior of APOBEC mutagenesis in IR regions was completely different. Figure 2a and Supplemental Figure S3 show that the density of APOBEC-induced mutations in IR genome regions increased with increasing APOBEC activity in cancer samples faster than the density of APOBEC mutagenesis in B-DNA genome regions. The effects observed for five of the considered non-canonical DNA structure types were statistically significant (Figure 2b). In the remaining APR motif, we observed no enrichment of the APOBEC mutagenesis compared to B-DNA genome regions at all levels of activity of APOBEC mutagenesis in cancer samples (Supplemental Figure S3). To assess the observed differences between the APOBEC mutation density in the non-canonical DNA structures and in B-DNA across the cancer samples, we calculated and visualized these densities in each sample (Figure 2c and Supplemental Figure S5).

We also compared the distribution of APOBEC-induced mutation clusters^28^ in non-B and B-DNA genome regions and did not find any statistically significant differences (data not shown). Additionally, the total size of APOBEC-enriched mutation clusters in a cancer sample could serve as an estimate of the fraction of hypermutable ssDNA in the genome formed during repair of double-strand breaks (DSB)^28^. Thus, we analyzed the dependence of the density of APOBEC-induced mutations in non-B genome regions on the total size of APOBEC-enriched mutation clusters. As the total cluster size increased, we observed the same trends as described above: relative decrease of APOBEC-induced mutation density in non-B DNA structures to the level in B-DNA and even below for DR, STR, MR, G4, and increase of the APOBEC-induced mutation density in IR (Supplemental Figure S4).

### Observed effects is not the result of the heterogeneity of APOBEC mutagenesis and non-B DNA structures along the genome

Then, we verified that the observed effects did not result from the known heterogeneity of APOBEC-induced mutations along the replication timing^12^, that is, the increased density of APOBEC-induced mutations in early-replicating regions and the decreased density in late-replicated regions. First, we calculated the distribution of APOBEC targets (TpC motifs) contained in the considered non-B DNA structures along the replication timing (Figure 3a). All types of non-B DNA structures, except for A-phased repeats, showed either an almost uniform distribution (MR and IR) or a distribution skewed towards increased fraction of TpC motifs, residing in non-canonical DNA structures, in early-replicating genome regions (DR, STR, G4). In contrast, TpC motifs contained in A-phased repeats were enriched in late-replicating regions.

**Figure 3.**
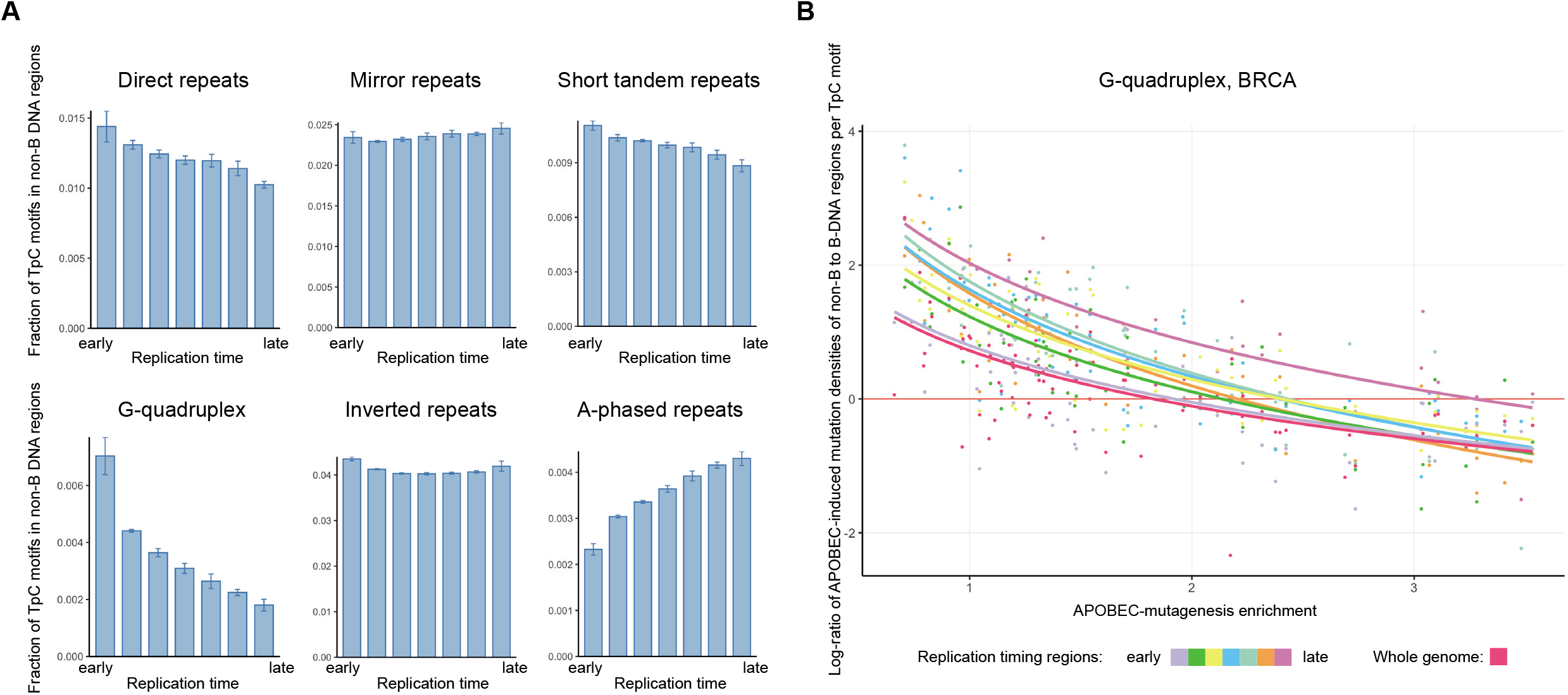
(a) Distribution of the non-B DNA structures along the replication timing. (b) Dependence of the log-ratio of APOBEC-induced mutation densities in G-quadruplex to B-DNA genome regions on the activity of APOBEC mutagenesis in different replication timing bins for the BRCA cancer.

If the effects for DR, STR, MR, and G4, described in the previous section, were the result of APOBEC mutagenesis heterogeneity relative to the replication timing, we should expect a reduction of the number of non-B structures in the regions of enriched APOBEC mutagenesis, i.e., in early-replicating genome regions. However, we did not observe that for these four types of non-B DNA structures. For additional evidence, we calculated the APOBEC-signature mutation density in non-canonical DNA genome regions dividing the genome into seven separate replication timing bins, from the early to the late replication timing. Figure 3b shows the results for G-quadruplexes, which have the most skewed distribution toward early-replicating regions, in the BRCA cancer, the cancer type having the largest number of samples. In this example and for the majority of other non-B DNA structures (data not shown), the observed effect of relative decrease of the APOBEC mutagenesis in most non-B regions as the APOBEC activity increases, is visible for each replication bin, i.e., in all sets of genome regions with approximately same replication timing. Thus, we conclude that the observed effects are not a consequence of the mutational heterogeneity of the APOBEC mutagenesis along the replication timing.

### Increase of APOBEC mutagenesis in inverted repeats, apparently associated with DNA secondary structure

To understand possible causes of the observed characteristics of APOBEC mutagenesis in non-B DNA genome regions, we analyzed the distribution of APOBEC-induced mutations relative to the secondary structure of non-B DNA motifs. We first analyzed the distribution of APOBEC-induced mutations in IR motifs. IR genome regions form hairpin-loop arms on both DNA strands. We stratified detected APOBEC-induced mutations into the ones that occurring in the hairpin stem and in the hairpin loop. Following recent reports on the propensity of APOBEC enzymes toward cytosine located at the 3’-end of the hairpin loop ^16,29^ we also further divided the hairpin loop mutations into two categories, ones occurring in the cytosine at the 3’-end of a loop (here and further 3’LEC (3’-loop end cytosine)) and all other positions of the hairpin loop (Figure 4a).

**Figure 4.**
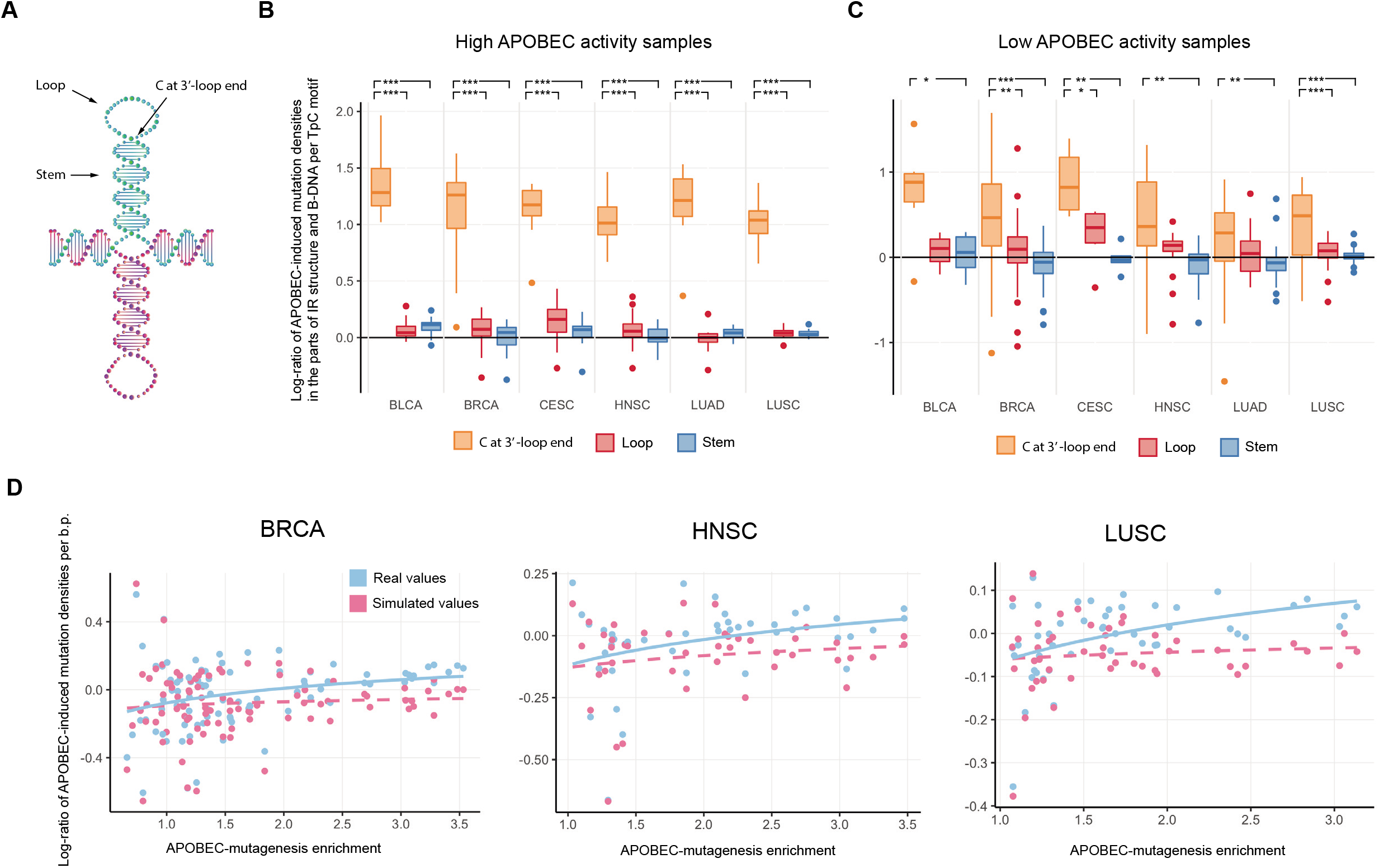
(a) Different parts of the IR motif’s secondary structure: stem, loop, and the cytosine at the 3’-end of the hairpin loop. (b,c) Distribution of APOBEC-induced mutation densities at TpC motifs in different parts of the IR secondary structure in cancer samples with high (b) and low (c) APOBEC activity. (d) Contribution of mutations at the cytosine at the 3’-end of the IR hairpin loop to the overall IR motif APOBEC-induced mutation load. Plots for different cancer types demonstrate the dependency of the APOBEC-induced mutation density per base pair on the activity of APOBEC mutagenesis for real and simulated data. In simulated data, the APOBEC-induced mutation density at the cytosine at the 3’-end of the IR hairpin loop was replaced by the average APOBEC-induced mutation density in the IR hairpin stem and the rest of the hairpin loop. Notation: *** – P-value < 0.001, ** – P-value < 0.01, * – P-value < 0.05.

Figure 4b shows that the density of APOBEC-induced mutations in IR genome regions in the 3’LECs is significantly higher than the density in the loops and stems observed in samples with high APOBEC activity (APOBEC enrichment > 2.0). Figure 4c shows that this effect is also less prominent but still observable in samples with low APOBEC activity (APOBEC enrichment < 2.0). Surprisingly, when we extended this analysis to APOBEC-negative cancer types (Supplemental Figure S6), we still observed this effect. This finding raises the question of whether the 3’LEC is a APOBEC-specific mutational hotspot.

To elucidate the impact of APOBEC-induced mutagenesis at the 3’-end of IR hairpin loops to the APOBEC-mutagenesis of the whole IR genome regions, we recalculated the mutation density by removing mutations at 3’LEC positions and replaced them by the average density of APOBEC-signature mutations in the IR hairpin loop and stem. Figure 4d shows that the increased density of APOBEC-signature mutagenesis at 3’LEC significantly impacts the accumulation of APOBEC-induced mutations in IR genome regions (P-values of the linear regression coefficients < 0.05 for four out of six cancer types). Thus, if the density of APOBEC-induced mutations at 3’LEC were not increased, then the overall density of APOBEC mutagenesis in IR regions would be approximately the same as in B-DNA regions (coefficients of the linear regression are not significantly different from zero), while actually it increases in comparison with the APOBEC-induced density in B-DNA with increasing of APOBEC activity.

### Single-stranded DNA complementary to the G-quadruplex four-stranded structure is not a favorable target of APOBEC enzymes

We also compared the frequency of APOBEC-induced mutations in G-quadruplex genome regions in the DNA strand with a four-stranded guanine-rich structure and in the complementary, unstructured DNA strand. Firstly, we calculated the density of TpC motifs in these two strands (Figure 5a). As expected, the DNA strand of G4 with a four-stranded structure showed a reduced number of TpC motifs in comparison with the average density of TpC motifs in B-DNA genome regions, as a larger fraction of the nucleotide sequence was guanines. On the other hand, the density of TpC motifs on the opposite strand was almost two-fold higher that the average density of TpC motifs in B-DNA. This increase was obviously due to stretches of cytosines complementary to guanines in the guanine-rich strand.

**Figure 5.**
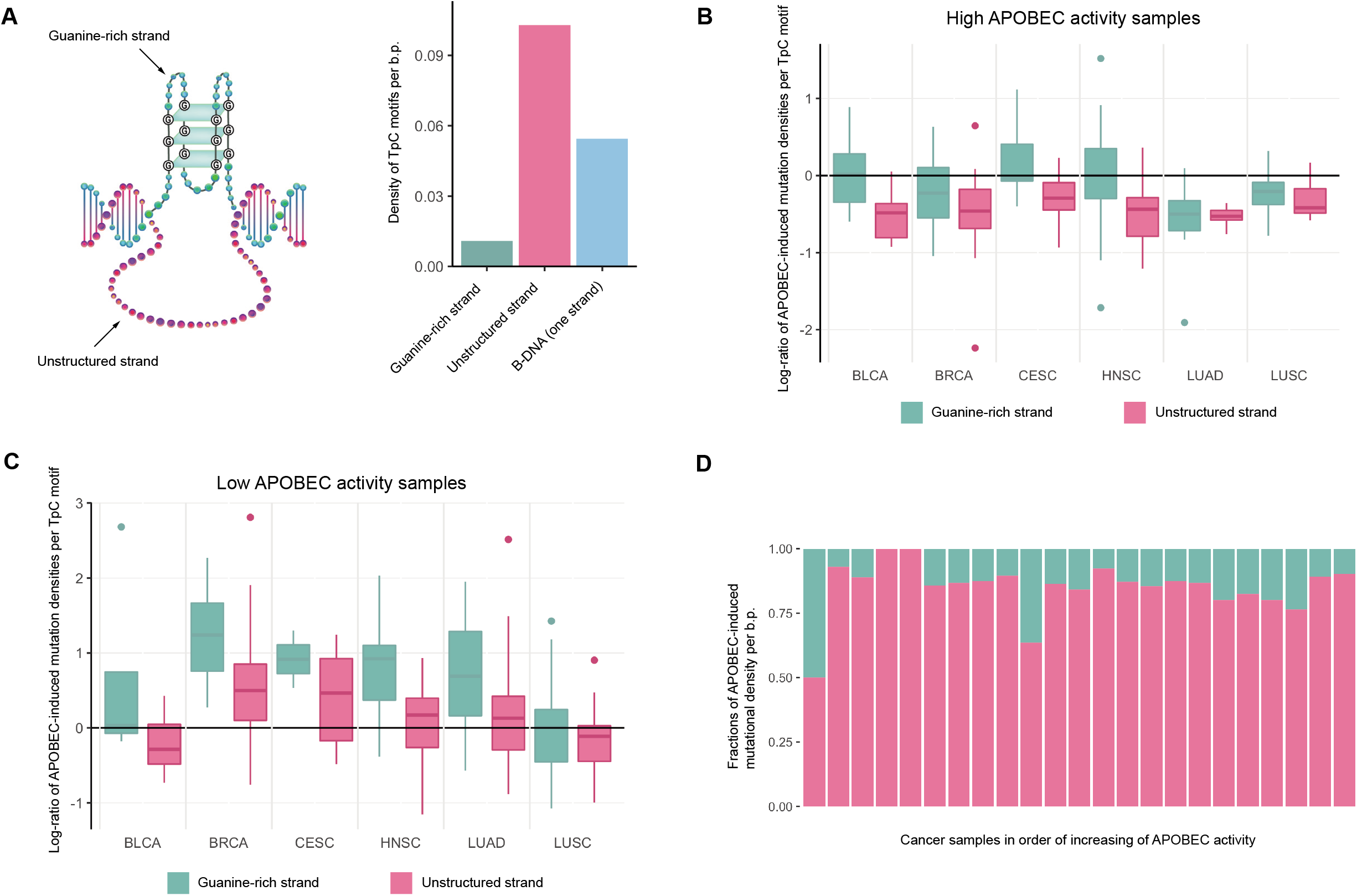
(a) Comparison of the densities of TpC motifs per base pair on both strands of G-quadruplex structures and B-DNA. (b,c) Distribution of APOBEC-induced mutational densities at TpC motifs on both strands of G-quadruplex structures in cancer samples with high (b) and low (c) APOBEC activity. (d) Impact of both strands of G-quadruplex structures on the APOBEC-induced mutation density per base pair in G4 genome regions.

Then, we calculated the densities of APOBEC-induced mutations in TpC motifs in both G4-motif strands. Surprisingly, the density of APOBEC-induced mutations in TpC motifs in the unstructured DNA strand of G4, which seemed to be a favorable APOBEC substrate, turned out to be smaller than the density in the guanine-rich strand. Notably, these difference in densities were observed in samples with both high activity of APOBEC mutagenesis (APOBEC enrichment > 2.0, Figure 5b) and low activity (APOBEC enrichment < 2.0, Figure 5c), and hence possibly represent an effect independent from the activity of the APOBEC mutagenesis. Moreover, in samples with high APOBEC activity, the density of APOBEC-induced mutations in the unstructured DNA strand of G4 was constantly smaller than the density in canonical double-strand DNA (Figure 5b,c), which also showed that this single-stranded DNA was not an optimal substrate of the APOBEC enzymes. At the same time, due to a small number of TpC motifs in the guanine-rich strand, its impact to the overall APOBEC-induced mutational density per base pair in G4 genome regions is small in comparison with the impact of the unstructured DNA strand of G4 (Figure 5e).

### Purine-pyrimidine alternations in Z-DNA lead to the absence of the APOBEC- and UV-mutageneses in this DNA structure

As described above, we observed a depletion of TpC motifs in Z-DNA genome regions (Supplemental Figure S2a) apparently due to the purine-pyrimidine alternations specific to the Z-DNA sequence^23,30^. The latter leads to a nearly complete absence of the APOBEC-mutagenesis in Z-DNA genome regions. While analyzing the density of somatic mutations in non-B regions in human cancers (Figure 1), we also found a prominent absence of the enrichment of somatic mutations in Z-DNA of skin cancers (MELA, SKCM). Using the UV-mutagenesis mutational signature, we calculated the density of UV-target motifs (TpC, CpC) and UV-associated C→T mutations. Similar to the APOBEC mutagenesis, we found that the apparent reason for the absence of UV-associated mutagenesis in Z-DNA is a depletion of UV-signature targets (Supplemental Figure S2b). Indeed, the fractions of TpC and CpC di-nucleotides in Z-DNA genome regions were 14- and 7-fold smaller than the fractions of the same di-nucleotides in B-DNA, respectively. Moreover, the density of UV-associated mutations per UV-target motifs was also decreased in Z-DNA in comparison with B-DNA (Supplemental Figure S2c, *P_MELA, TC_* = 2.2×10^−10^, *P_MELA, CC_* = 2.1×10^−15^, *P_SKCM, TC_* = 3.7×10^−5^, *P_SKCM, TC_* = 1.5×10^−7^).

## Discussion

Rapid accumulation of mutation data for cancer genomes provides an excellent opportunity to study mutational processes in human cancer and their heterogeneity along the genome. Understanding the mutational heterogeneity along the genome is an essential component in computational methods for the identification of cancer-associated genes^31^. It has been found earlier that genome regions with non-canonical DNA structure such as G-quadruplex, cruciform, various types of repeats, and Z-DNA, which cover together about 10% of the human genome, usually have an increased density of somatic^24^ and germline^27^ mutations. At the same time, classification of mutations into mutagen-related classes based on their nucleotide context using so-called mutational signatures in many cases allows for deciphering the individual impact of endogenous and exogenous mutagens. Here, we analyzed the activity of mutagenesis induced by APOBEC enzymes in non-B DNA genome regions, compared its level with the APOBEC mutagenesis in B-DNA, and elucidated the impact of structural parts of non-B DNA structures, focusing on ssDNA regions.

Using the large PCAWG dataset of mutations in cancer genomes, we confirmed that, in general, the density of somatic mutations is relatively larger in non-B DNA genome regions, as was found in other datasets of cancer mutagenesis^24,32^. With most of the mutagenic sources, mutations in cancer genomes originate from either misincorporation during DNA copying or from unrepaired lesions in dsDNA. Within this paradigm, the increased density of mutations in non-B DNA could be explained by a higher rate of replication errors or/and by lower access to error-prone DNA repair systems. We unexpectedly found that for four out of seven considered non-B DNA structures (DR, STR, MR, G4), the density of APOBEC-signature mutations decreases to the level in B-DNA as the APOBEC activity in cancer samples increases. Moreover, in samples with very high activity of APOBEC mutagenesis, the APOBEC mutational density in these non-B DNA structures was even lower than the density in B-DNA genome regions.

We speculate that the observed tendency to a similarity in APOBEC-induced mutation densities in B- and non-B DNA genome regions can be explained by the unique association of APOBEC mutagenesis with ssDNA, which can be generated during replication, double-strand break repair, and transcription^10^. Once APOBEC deaminates a cytosine in ssDNA, it can be accurately returned to non-mutant sequence by base-excision repair if the complementary strand is available as a repair template. However, if ssDNA is persistent, i.e., does not have access to the complementary template for accurate repair, the deaminated cytosine will be fixed into mutation in the next rounds of replication^10^. We propose that both non-B DNA and B-DNA sequences are equally prone to generating persistent ssDNA. In the case of non-B DNA structures, unwinding could be aided by special helicases^33^.

Moreover, a lower density of APOBEC mutations in non-B-DNA as compared to B-DNA regions that we observed in cancer samples with very high levels of APOBEC mutagenesis can also be explained in connection with the requirement of persistent ssDNA substrate for mutagenesis if persistent ssDNA formed in non-B DNA regions is less accessible by APOBEC deaminases compared to ssDNA formed in B-DNA. It is well established that non-B DNA structures cause replication stalling and require special systems aiding to replicate these regions^34^. This could lead to higher recruitment of replication protein A (RPA), which binds to ssDNA and is known to counteract APOBEC deamination^35,36^. An alternative or an additional explanation could be recruiting for replicating of non-B DNA regions the specific DNA polymerase called Primase-Polymerase (PrimPol). This polymerase is known to shield DNA from APOBEC/AID mutagenesis^37^. PrimPol is known for its implication in eukaryotic DNA damage tolerance^38^, displaying both translesion synthesis and (re)-priming properties^39^. PrimPol is required for replicating G-quadruplexes^40^ and presumably other types of non-canonical DNA structures^41^. It has been shown that PrimPol prevents mutagenesis of abasic sites induced by APOBEC/AID by repriming downstream of AP-sites on the leading strand, prohibiting error-prone TLS, and simultaneously stimulating error-free homology-directed repair^37^.

Despite this observed general trend, we found the opposite effect for the APOBEC mutagenesis in IR genome regions – the density of APOBEC-induced mutations increased with increasing APOBEC activity in cancer samples. We showed that a substantial part of this increase could be attributed to the mutagenesis in cytosine at the 3’-end of the IR hairpin loop, which was recently identified as the APOBEC’s hotspot^29^. However, our analysis of APOBEC-negative cancers indicates that this mutational hotspot might be not APOBEC-specific. We speculate that the increase of APOBEC mutagenesis in IR regions may be associated with double-strand breaks^1^ induced by the replication stalling at cruciform DNA structures^42^, as it is known that 5’ ends of these breaks are resected to generate 3’-protruding ssDNA regions for subsequent repair process^43^.

Apart from considering non-B DNA regions as a whole, we also analyzed when possible the distribution of mutation densities between the dsDNA and ssDNA parts of non-canonical DNA structures. Thus, in the cruciform structure (IR motif), the cytosine at the 3’-end of the IR hairpin loop showed the largest APOBEC-induced mutational density in comparison with the densities in the IR hairpin stem and other parts of the loop. We also focused on the large stretches of presumably single-stranded DNA opposite to the G-quadruplex's guanine-rich strand. Surprisingly, we found that APOBEC-induced mutational density per TpC motif in this ssDNA is less than the density both in the guanine-rich strand of the G-quadruplex and in B-DNA. We suggest that a reason for that could be occupation of this ssDNA by other single strand-binding proteins^44,45^ or formation of cell cycle-depending alternative secondary structure from the stacked cytosines, called the intercalated motif (i-motif)^46^. Additionally, we verified that ssDNA genome regions detected in vivo^47^ actually intersected with computationally predicted G4 regions used in this study (Supplemental Figure S7).

The analysis of APOBEC-induced mutation density in A-phased repeats did not show any statistically significant differences in comparison with B-DNA. Z-DNA was excluded from the analysis as we found a negligible number of TpC motifs in these genome regions, as Z-DNA is formed by purine-pyrimidine alternating sequences. Interestingly, we have observed that this also decreases the relative level of UV-mutagenesis in Z-DNA genome regions and hence provides one more example of reduced cancer mutagenesis in non-B DNA genome regions.

Overall, we have observed unexpectedly low densities of APOBEC-induced somatic mutations in non-canonical DNA structures in human cancers, which, in contrast to mutations caused by other mutagens, is approximately equal or even smaller than the level of APOBEC-mutagenesis in B-DNA, despite the presence of ssDNA in most of non-B DNA structures. Elucidation of the mechanistic basis of these observations requires further research.

## Methods

Somatic mutations were taken from the `Pan-Cancer Analysis of Whole Genomes` project^25^. Six cancer types, BLCA, BRCA, HNSC, LUAD, LUSC, CESC, each having a substantial number of samples enriched with the APOBEC mutagenesis signature (APOBEC-mutagenesis enrichment > 2.0, calculated as in ref.^8^), were selected for the analysis. Non-B DNA regions were downloaded from `Non-B DB`^26^. Seven non-B DNA motifs were considered: G-quadruplex, inverted repeats, mirror repeats, direct repeats, A-phased repeats, short tandem repeats, and Z-DNA. Information on non-B DNA motif secondary structure was obtained from the `Non-B DB` annotations. Human genome assembly `GRCh37/hg19` was used as the reference. The mutation density D_APOBEC_ of the APOBEC mutagenesis per target in a particular genome region was calculated as the number of single-base substitutions <u>C</u>→T or <u>C</u>→G in the TpC motif, divided by the total number of the TpC motifs in this region: *D*_APOBEC_ = *N*_APOBEC_ / *N*_TCN_. The same density per base pair was calculated as the number of single-base substitutions <u>C</u>→T or <u>C</u>→G in the TpC motif divided by the genome region size. Statistical significance of the difference in the mutation density between genome regions was calculated by the 10000-fold random shuffling of mutation positions in each chromosome of each sample.

Single strand DNA-sequencing data was taken from ref.^47^. Short reads were aligned to the human (`hg19`) genome using `Bowtie2` (ver. 2.3.0). Only unique mappings were kept using `samtools`^48^. Sequence reads with a mapping quality of less than 30 were filtered out. SAM-file was converted to the `BEDGRAPH` format using `bedtools`^49^. Data on APOBEC-induced mutational clusters was taken from ref.^28^. Coordinates of CpG islands were taken from the `UCSC Genome Browser`^50^.

## Acknowledgements

We thank Dmitry Gordenin for thoughtful discussion, critical reading and editing of the manuscript and Irina Ponomareva for the non-B DNA structures artworks. This study was partially supported by RFBR (grant 18-29-13011) to M.S.G. and by InterSystems via Innovations Program grant to M.D.K.. The authors would like to acknowledge the dbGaP repository for providing access to the TCGA dataset (the accession number is phs000178.v11.p8).

## Author Contributions

M.D.K. conceived the study. G.V.P., B.F., V.A.N., R.A., E.S., N.L., A.A.D, and A.A.C. performed the calculations. All authors contributed to data analysis. M.S.G. and M.D.K. wrote the paper.

## Notes

### Competing Interest Statement

The authors have declared no competing interest.

## References

1. Roberts, S. A. et al. Clustered Mutations in Yeast and in Human Cancers Can Arise from Damaged Long Single-Strand DNA Regions. Mol. Cell 46, 424–435 (2012).

2. Nik-Zainal, S. et al. Mutational processes molding the genomes of 21 breast cancers. Cell 149, 979–993 (2012).

3. Burns, M. B. et al. APOBEC3B is an enzymatic source of mutation in breast cancer. Nature 494, 366–370 (2013).

4. Salter, J. D., Bennett, R. P. & Smith, H. C. The APOBEC Protein Family: United by Structure, Divergent in Function. Trends Biochem. Sci. 41, 578–594 (2016).

5. Shi, K. et al. Structural basis for targeted DNA cytosine deamination and mutagenesis by APOBEC3A and APOBEC3B. Nat. Struct. Mol. Biol. 24, 131–139 (2017).

6. Alexandrov, L. B. et al. Signatures of mutational processes in human cancer. Nature 500, 415–421 (2013).

7. Burns, M. B., Temiz, N. A. & Harris, R. S. Evidence for APOBEC3B mutagenesis in multiple human cancers. Nat. Genet. 45, 977–983 (2013).

8. Roberts, S. A. et al. An APOBEC cytidine deaminase mutagenesis pattern is widespread in human cancers. Nat. Genet. 45, 970–976 (2013).

9. Yang, B., Li, X., Lei, L. & Chen, J. APOBEC: From mutator to editor. J. Genet. Genomics 44, 423–437 (2017).

10. Saini, N. & Gordenin, D. A. Hypermutation in single-stranded DNA. DNA Repair (Amst). 91–92, 102868 (2020).

11. Haradhvala, N. J. et al. Mutational Strand Asymmetries in Cancer Genomes Reveal Mechanisms of DNA Damage and Repair. Cell 164, 538–549 (2016).

12. Kazanov, M. D. et al. APOBEC-Induced Cancer Mutations Are Uniquely Enriched in Early-Replicating, Gene-Dense, and Active Chromatin Regions. Cell Rep. 13, 1103–1109 (2015).

13. Chervova, A. et al. Analysis of gene expression and mutation data points on contribution of transcription to the mutagenesis by APOBEC enzymes. NAR Cancer 3, 1–12 (2021).

14. Seplyarskiy, V. B. et al. APOBEC-induced mutations in human cancers are strongly enriched on the lagging DNA strand during replication. Genome Res. 26, 174–182 (2016).

15. Zhang, Y., Zhou, H. & Ou-Yang, Z. C. Stretching single-stranded DNA: Interplay of electrostatic, base-pairing, and base-pair stacking interactions. Biophys. J. 81, 1133–1143 (2001).

16. Langenbucher, A. et al. An extended APOBEC3A mutation signature in cancer. Nat. Commun. 12, 1–11 (2021).

17. Zhao, J., Bacolla, A., Wang, G. & Vasquez, K. M. Non-B DNA structure-induced genetic instability and evolution. Cell. Mol. Life Sci. 67, 43–62 (2010).

18. Gordenin, D. A. et al. Inverted DNA repeats: a source of eukaryotic genomic instability. Mol. Cell. Biol. 13, 5315–5322 (1993).

19. Brázda, V., Laister, R. C., Jagelská, E. B. & Arrowsmith, C. Cruciform structures are a common DNA feature important for regulating biological processes. BMC Mol. Biol. 12, (2011).

20. Frank-Kamenetskii, M. D. & Mirkin, S. M. Triplex DNA structures. Annu. Rev. Biochem. 64, 65–95 (1995).

21. Spiegel, J., Adhikari, S. & Balasubramanian, S. The Structure and Function of DNA G-Quadruplexes. Trends Chem. 2, 123–136 (2020).

22. Sinden, R. R., Pytlos-Sinden, M. J. & Potaman, V. N. Slipped strand DNA structures. Front. Biosci. 12, 4788–4799 (2007).

23. Ravichandran, S., Subramani, V. K. & Kim, K. K. Z-DNA in the genome: from structure to disease. Biophys. Rev. 383–387 (2019). doi:10.1007/s12551-019-00534-1

24. Georgakopoulos-Soares, I., Morganella, S., Jain, N., Hemberg, M. & Nik-Zainal, S. Noncanonical secondary structures arising from non-B DNA motifs are determinants of mutagenesis. Genome Res. 28, 1264–1271 (2018).

25. Campbell, P. J. et al. Pan-cancer analysis of whole genomes. Nature 578, 82–93 (2020).

26. Cer, R. Z. et al. Non-B DB v2.0: A database of predicted non-B DNA-forming motifs and its associated tools. Nucleic Acids Res. 41, 94–100 (2013).

27. Guiblet, W. M. et al. Non-B DNA: a major contributor to small- and large-scale variation in nucleotide substitution frequencies across the genome. Nucleic Acids Res. 49, 1497–1516 (2021).

28. Sakofsky, C. J. et al. Repair of multiple simultaneous double-strand breaks causes bursts of genome-wide clustered hypermutation. PLOS Biol. 17, e3000464 (2019).

29. Buisson, R. et al. Passenger hotspot mutations in cancer driven by APOBEC3A and mesoscale genomic features. Science (80-.). 364, (2019).

30. Rich, A. & Zhang, S. Timeline: Z-DNA: the long road to biological function. Nat. Rev. Genet. 4, 566–572 (2003).

31. Lawrence, M. S. et al. Mutational heterogeneity in cancer and the search for new cancer-associated genes. Nature 499, 214–218 (2013).

32. McKinney, J. A. et al. Distinct DNA repair pathways cause genomic instability at alternative DNA structures. Nat. Commun. 11, 1–12 (2020).

33. Sharma, S. Non-B DNA Secondary Structures and Their Resolution by RecQ Helicases. J. Nucleic Acids 2011, 724215 (2011).

34. Wang, G. & Vasquez, K. M. Effects of replication and transcription on DNA Structure-Related genetic instability. Genes (Basel). 8, (2017).

35. Wong, L., Vizeacoumar, F. S., Vizeacoumar, F. J. & Chelico, L. APOBEC1 cytosine deaminase activity on single-stranded DNA is suppressed by replication protein A. Nucleic Acids Res. 49, 322–339 (2021).

36. Brown, A. L. et al. Single-stranded DNA binding proteins influence APOBEC3A substrate preference. Sci. Rep. 11, 1–13 (2021).

37. Pilzecker, B. et al. PrimPol prevents APOBEC/AID family mediated DNA mutagenesis. Nucleic Acids Res. 44, 4734–4744 (2016).

38. Bailey, L. J., Bianchi, J. & Doherty, A. J. PrimPol is required for the maintenance of efficient nuclear and mitochondrial DNA replication in human cells. Nucleic Acids Res. 47, 4026–4038 (2019).

39. Bainbridge, L. J., Teague, R. & Doherty, A. J. Repriming DNA synthesis: An intrinsic restart pathway that maintains efficient genome replication. Nucleic Acids Res. 49, 4831–4847 (2021).

40. Schiavone, D. et al. PrimPol Is Required for Replicative Tolerance of G Quadruplexes in Vertebrate Cells. Mol. Cell 61, 161–169 (2016).

41. Šviković, S. et al. R-loop formation during S phase is restricted by PrimPol-mediated repriming. EMBO J. 38, 1–19 (2019).

42. Lu, S. et al. Short inverted repeats are hotspots for genetic instability: Relevance to cancer genomes. Cell Rep. 10, 1674–1680 (2015).

43. Ceccaldi, R., Rondinelli, B. & D’Andrea, A. D. Repair Pathway Choices and Consequences at the Double-Strand Break. Trends Cell Biol. 26, 52–64 (2016).

44. Lacroix, L. et al. Identification of two human nuclear proteins that recognise the cytosine-rich strand of human telomeres in vitro. Nucleic Acids Res. 28, 1564–1575 (2000).

45. Kang, H.-J., Kendrick, S., Hecht, S. M. & Hurley, L. H. The transcriptional complex between the BCL2 i-motif and hnRNP LL is a molecular switch for control of gene expression that can be modulated by small molecules. J. Am. Chem. Soc. 136, 4172–4185 (2014).

46. Abou Assi, H., Garavís, M., González, C. & Damha, M. J. i-Motif DNA: structural features and significance to cell biology. Nucleic Acids Res. 46, 8038–8056 (2018).

47. Kouzine, F. et al. Permanganate/S1 Nuclease Footprinting Reveals Non-B DNA Structures with Regulatory Potential across a Mammalian Genome. Cell Syst. 4, 344–356.e7 (2017).

48. Danecek, P. et al. Twelve years of SAMtools and BCFtools. Gigascience 10, (2021).

49. Quinlan, A. R. & Hall, I. M. BEDTools: a flexible suite of utilities for comparing genomic features. Bioinformatics 26, 841–842 (2010).

50. Kent, W. J. et al. The human genome browser at UCSC. Genome Res. 12, 996–1006 (2002).

