## Supplemental Figures for "APOBEC mutagenesis is low in most types of non-B DNA structures, unlike other types of cancer mutagenesis"

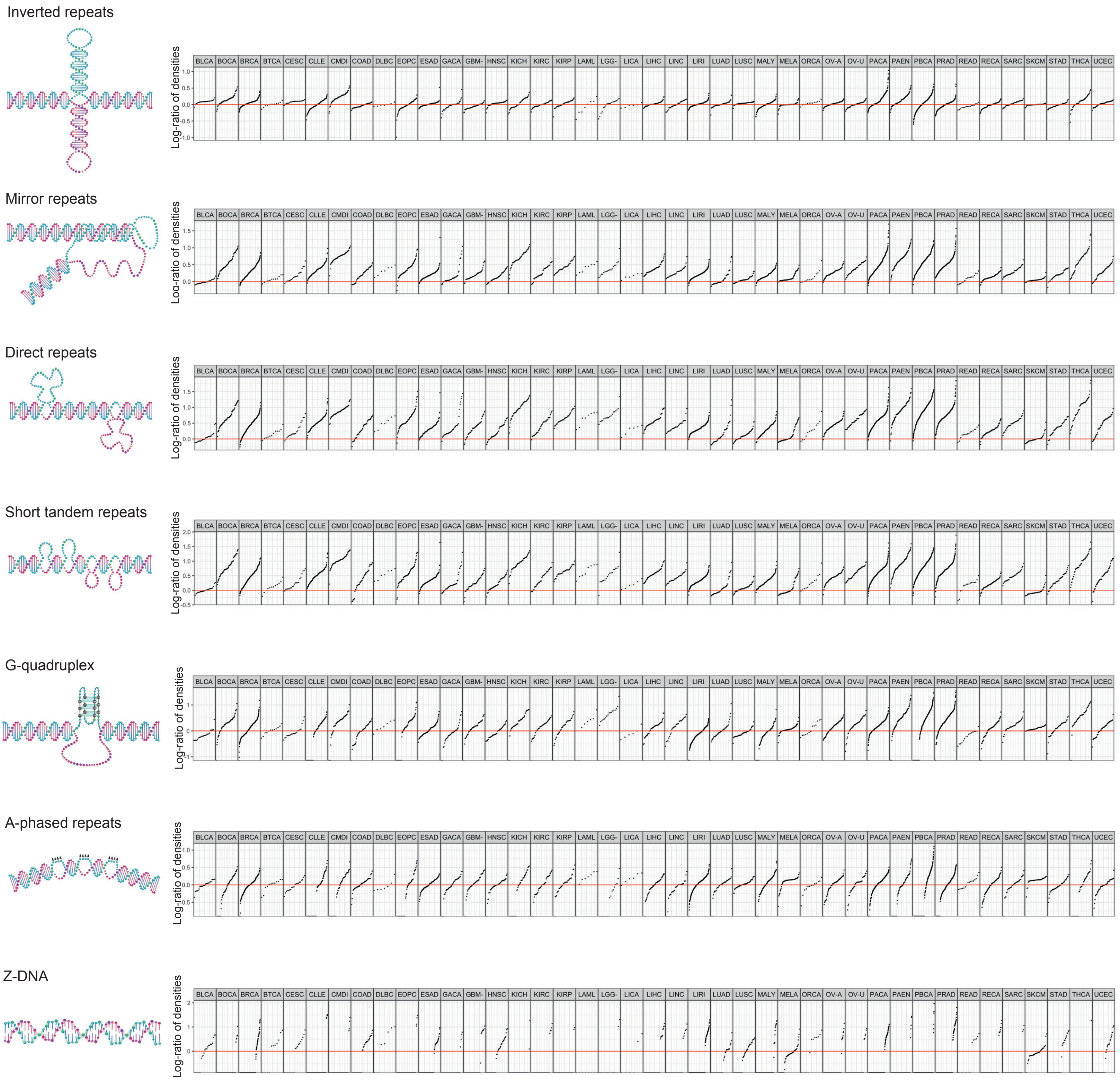


Figure S1. Enrichment of APOBEC-signature (TpC) somatic single-base substitutions (SBS) observed in various types of non-canonical DNA regions and different cancers. Single dot corresponds to a particular cancer sample, with the vertical position indicating the log-ratio of the mutational density in non-B DNA structure genome regions to the mutational density in B-DNA genome regions.

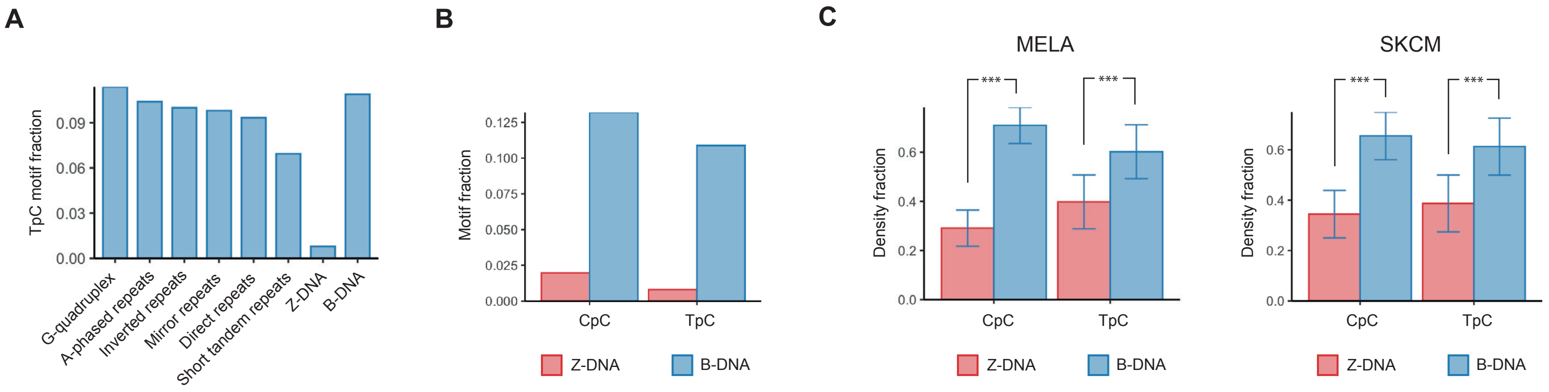


Figure S2. (a) Fraction of the APOBEC-signature motif, TpC, in each type of non-B DNA structure. (b) Fraction of the UV-signature motifs, CpC and TpC, in Z- and B-DNA. (c) Densities of UV-induced mutations in TpC and CpC di-nucleotides in Z- and B-DNA. Notation: *** – *P*-value < 0.001.

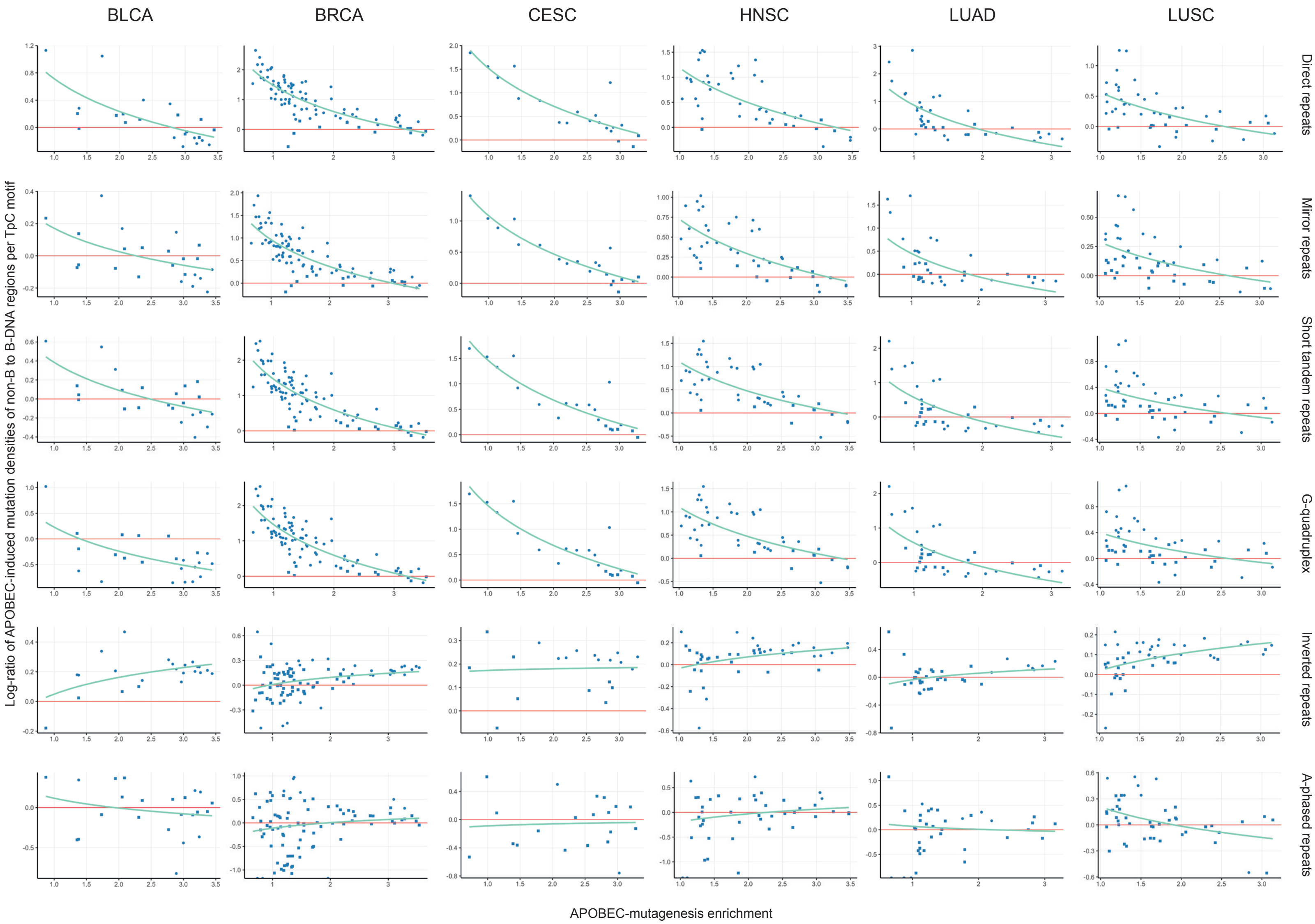


Figure S3. Dependence of the log-ratio of APOBEC-induced mutation densities in non-B DNA to B-DNA genome regions on the activity of APOBEC mutagenesis in cancer samples. Point shape – round or square – corresponds to significant or insignificant statistical differences between two densities in a particular cancer sample, respectively.

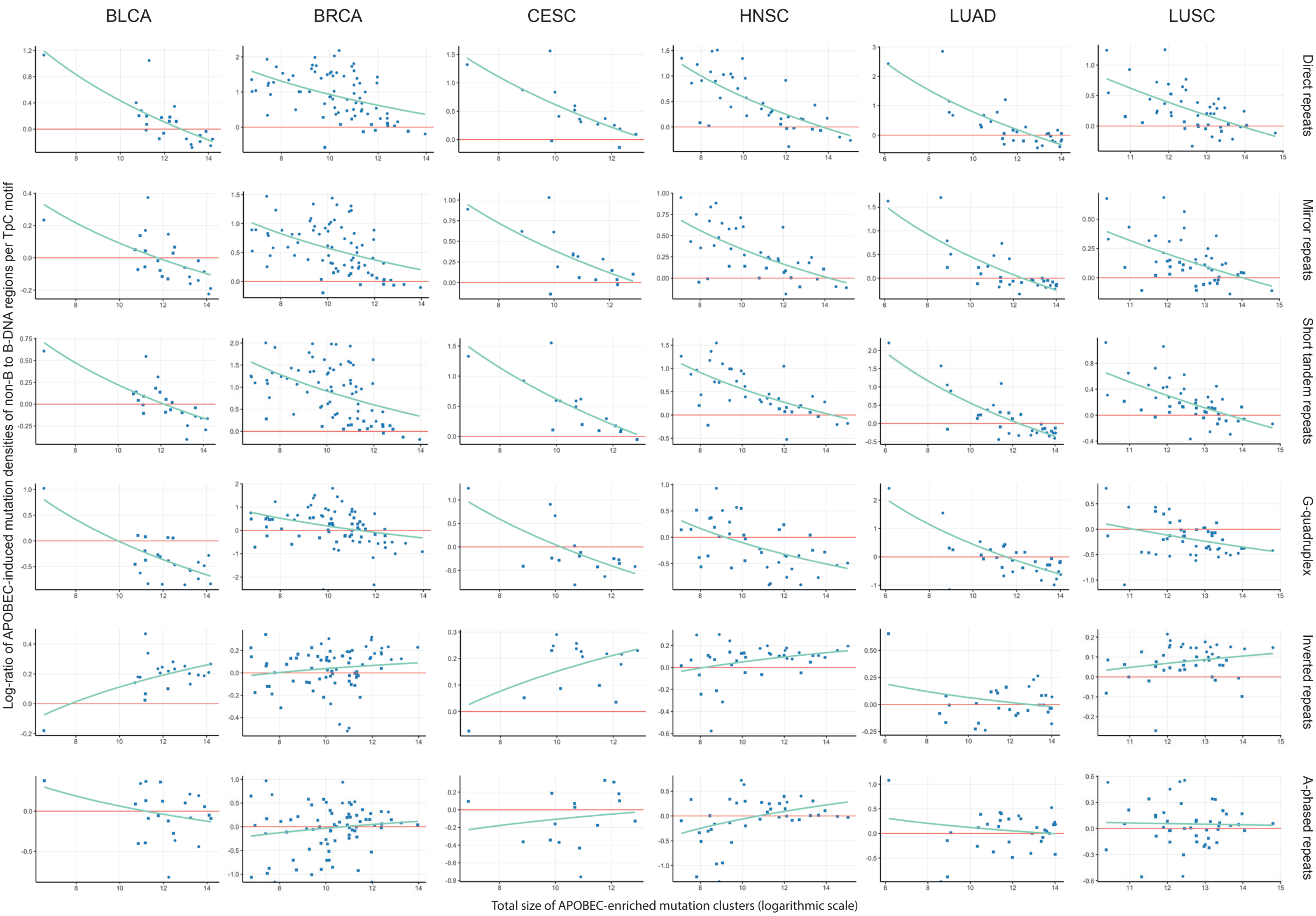


Figure S4. Dependence of the log-ratio of APOBEC-induced mutation densities in non-B DNA to B-DNA genome regions on the total size of APOBEC-enriched mutation clusters in cancer samples. Notation as in Fig. S3.

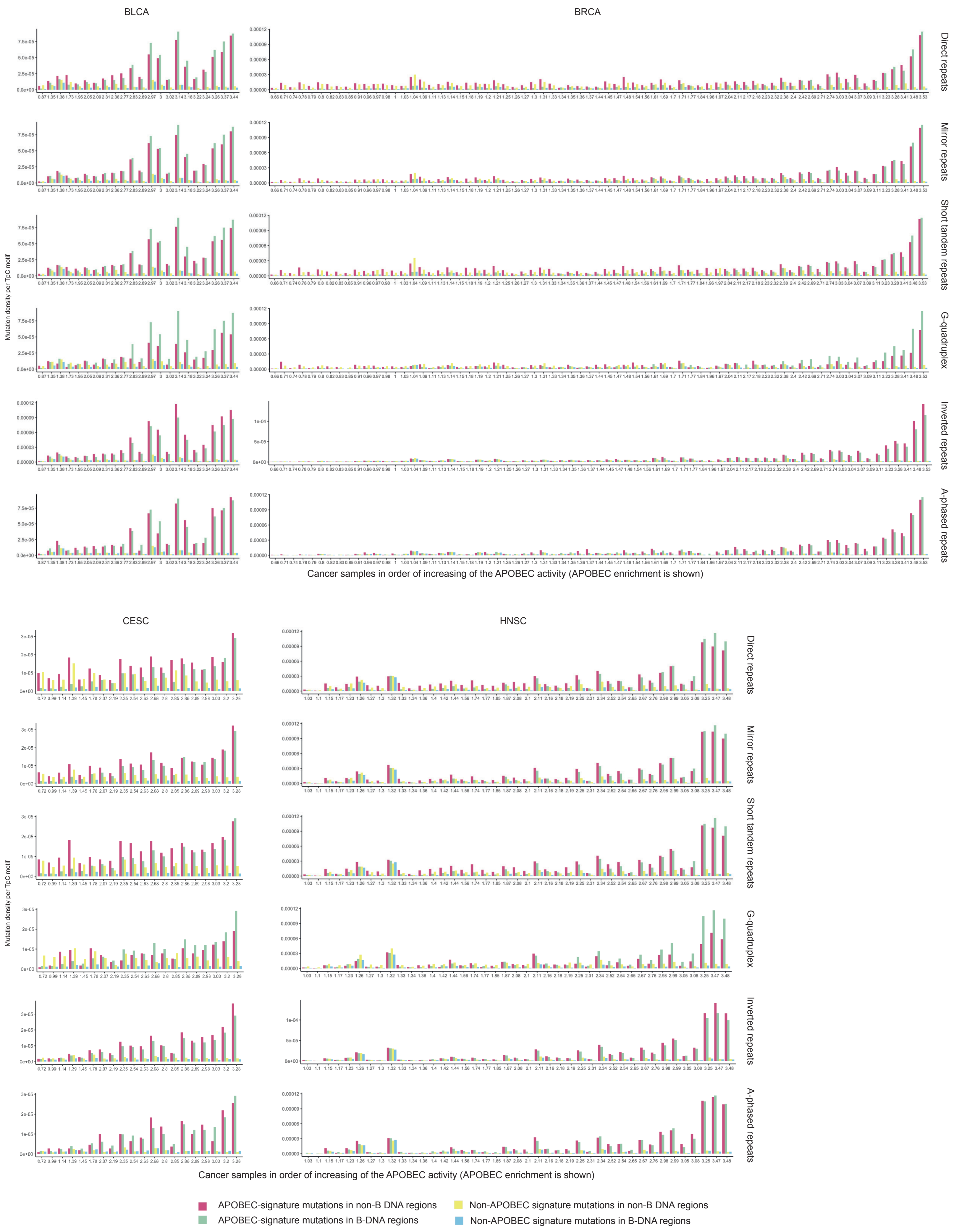
BLCA
BRCA
CESC
HNSC
Direct repeats
Mirror repeats
Short tandem repeats
G-quadruplex
Inverted repeats
A-phased repeats
Mutation density per TpC motif
Cancer samples in order of increasing of the APOBEC activity (APOBEC enrichment is shown)
APOBEC-signature mutations in non-B DNA regions
APOBEC-signature mutations in B-DNA regions
Non-APOBEC signature mutations in non-B DNA regions
Non-APOBEC signature mutations in B-DNA regions

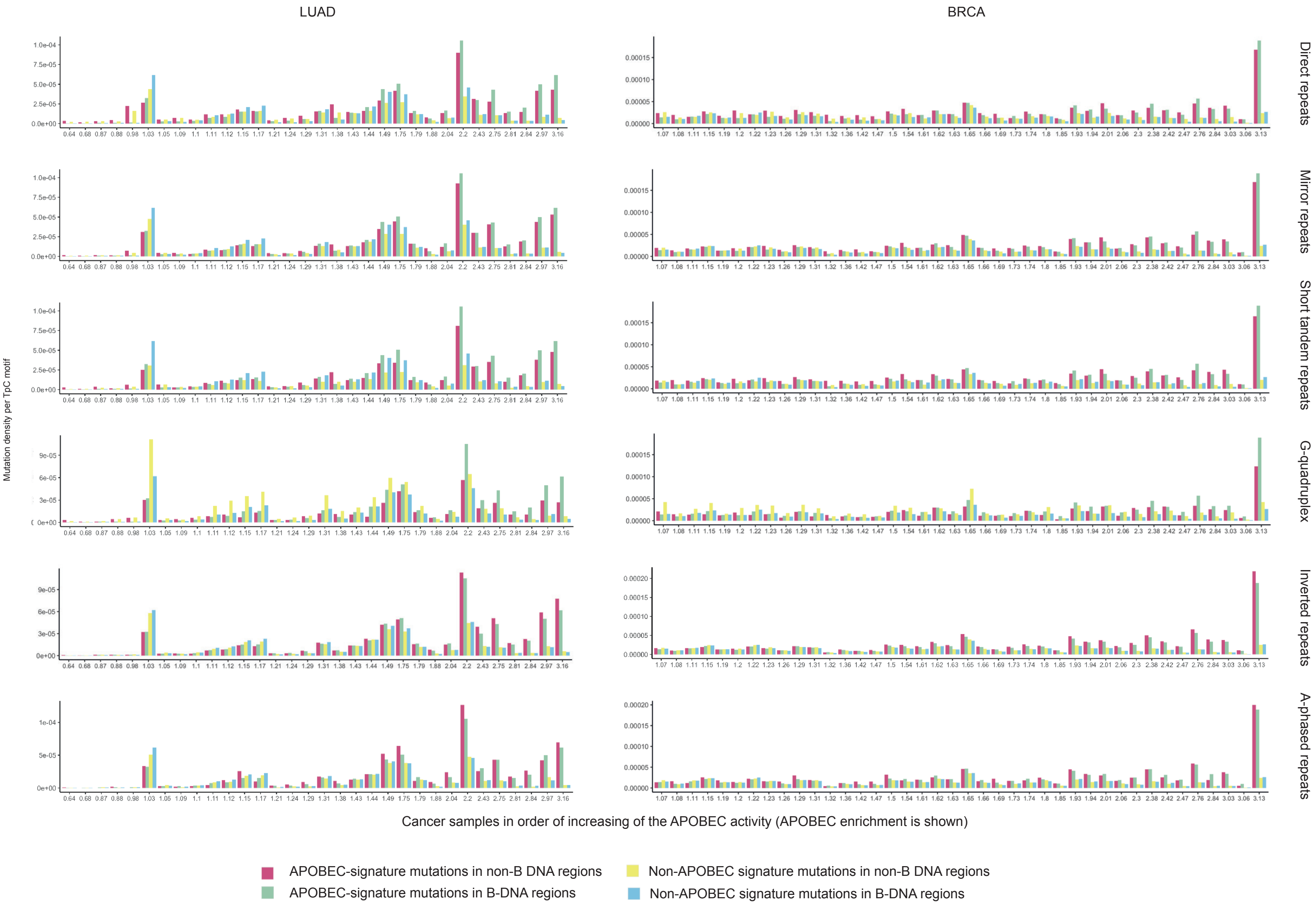


Figure S5. APOBEC-induced and other mutation densities in non-B and B-DNA genome regions.

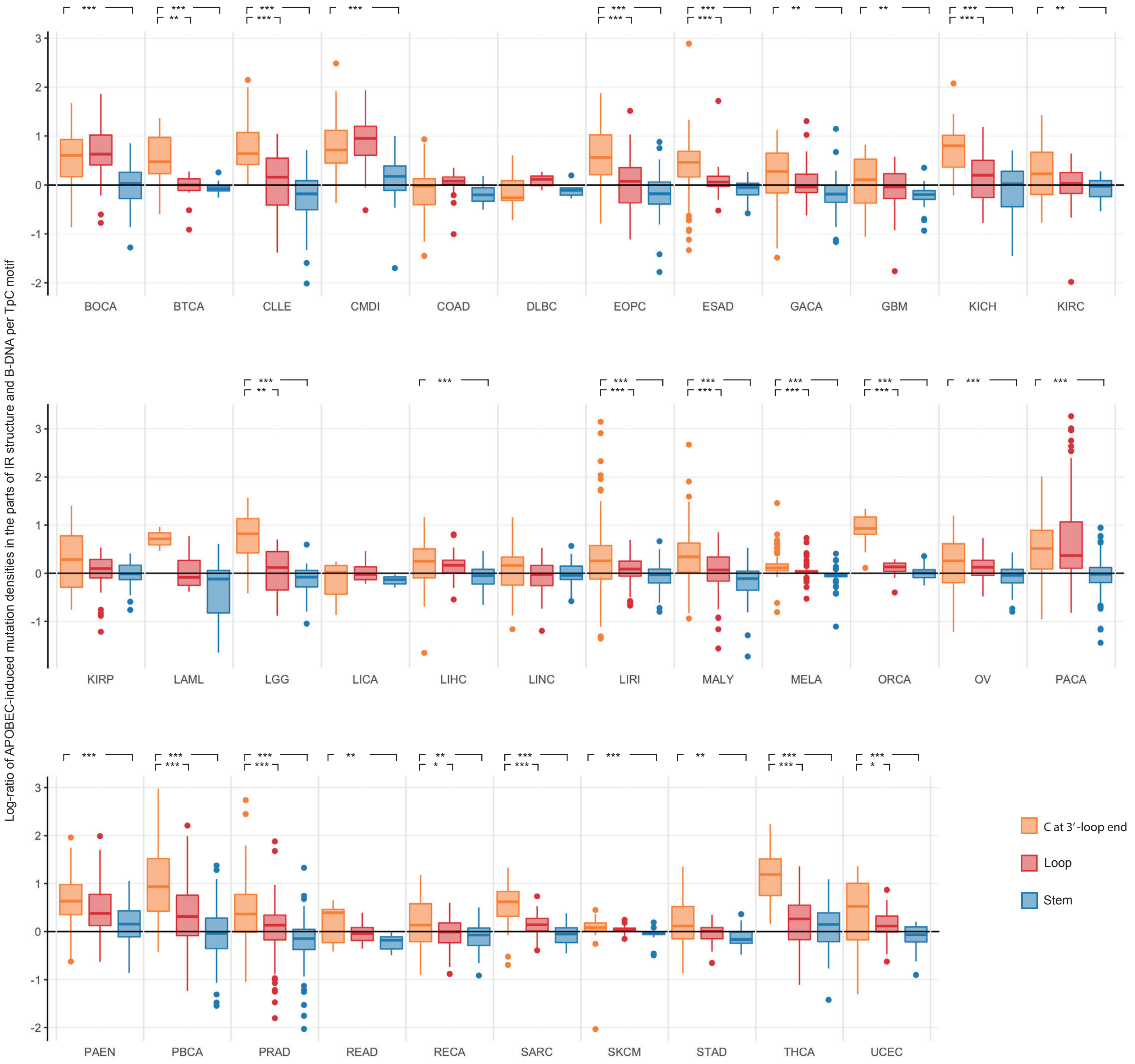


Figure S6. Distribution of APOBEC-signature mutational densities at TpC motifs in different parts of the IR secondary structure in APOBEC-negative cancers. Notation: *** – *P*-value < 0.001, ** – *P* -value < 0.01, * – *P* -value < 0.05.

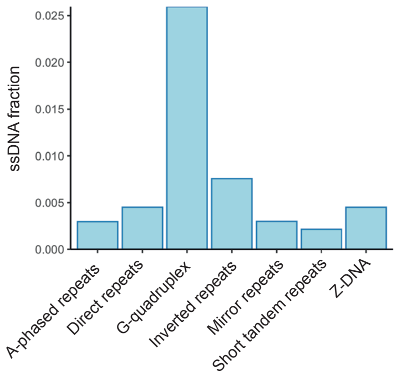


Figure S7. Fraction of ssDNA in non-B DNA structures.
